# Comparison of large-scale citizen science data and long-term study data for phenology modeling

**DOI:** 10.1101/335802

**Authors:** Shawn D. Taylor, Joan M. Meiners, Kristina Riemer, Michael C. Orr, Ethan P. White

## Abstract

Large-scale observational data from citizen science efforts are becoming increasingly common in ecology, and researchers often choose between these and data from intensive local-scale studies for their analyses. This choice has potential trade-offs related to spatial scale, observer variance, and inter-annual variability. Here we explored this issue with phenology by comparing models built using data from the large-scale, citizen science National Phenology Network (NPN) effort with models built using data from more intensive studies at Long Term Ecological Research (LTER) sites. We built process based phenology models for species common to each dataset. From these models we compared parameter estimates, estimates of phenological events, and out-of-sample errors between models derived from both NPN and LTER data. We found that model parameter estimates for the same species were most similar between the two datasets when using simple models, but parameter estimates varied widely as model complexity increased. Despite this, estimates for the date of phenological events and out-of-sample errors were similar, regardless of the model chosen. Predictions for NPN data had the lowest error when using models built from the NPN data, while LTER predictions were best made using LTER-derived models, confirming that models perform best when applied at the same scale they were built. Accordingly, the choice of dataset depends on the research question. Inferences about species-specific phenological requirements are best made with LTER data, and if NPN or similar data are all that is available, then analyses should be limited to simple models. Large-scale predictive modeling is best done with the larger-scale NPN data, which has high spatial representation and a large regional species pool. LTER datasets, on the other hand, have high site fidelity and thus characterize inter-annual variability extremely well. Future research aimed at forecasting phenology events for particular species over larger scales should develop models which integrate the strengths of both datasets.

## Introduction

Plant phenology, the timing of recurring biological events such as flowering, plays an important role in ecological research extending from local to global scales (Cleland et al., 2007; Richardson et al., 2013; Tang et al., 2016). At large scales, uncertainty in the timing of spring leaf out and fall senescence influence the carbon budget of earth system models, which has implications for correctly accounting for biosphere-atmosphere feedbacks in long-term climate forecasts (Richardson et al., 2012). At smaller scales, species-specific responses to temperature and precipitation can alter flower communities (Diez et al., 2012; CaraDonna et al., 2014; Theobald et al., 2017) and affect the abundance and richness of both pollinators (Ogilvie and Forrest, 2017; Ogilvie et al., 2017) and organisms at higher trophic levels (Tylianakis et al., 2008). Plant phenology models that are robust at multiple ecological scales, or deemed appropriate for a particular scale, are needed to better understand and forecast the timing of key biological events.

Many plant phenology studies use intensively collected datasets from a single location over a long time-period by a single research group (Cook et al., 2012; Wolkovich et al., 2012; Iler et al., 2013; Roberts et al., 2015). These datasets have regular sampling and large numbers of samples over long periods of time. As a result, the biological and climatic variability at that site is well represented. It is common for phenology models built with observations from a single site to not transfer well to other sites (García-Mozo et al., 2008; Xu and Chen, 2013; Olsson and Jönsson, 2014; Basler, 2016). This lack of transferability can be driven by plasticity in phenology requirements, local adaptation, microclimates, or differences in plant age or population density (Kramer, 1995; Diez et al., 2012). For these reasons, data from a single location is not adequate for larger scale phenology modeling. Accurately forecasting phenology at larger scales will require models that account for the full range of variation across a species’ range (Richardson et al., 2013; Tang et al., 2016; Chuine and Régnière, 2017), which will necessitate the use of data sources beyond traditional single-site studies.

Data from citizen science projects is becoming increasingly important for ecological research (Kelling et al., 2009; Dickinson et al., 2010; Tulloch et al., 2013). Because this data is often collected by large numbers of volunteers, it is possible to gather data at much larger scales than with individual research teams. A relatively new citizen science project started in 2009, The National Phenology Network (NPN), collects phenology observations from volunteers throughout North America and makes the data openly available (Schwartz et al., 2012). Data from this project has already been used to study variation in oak phenology at a continental scale (Gerst et al., 2017), develop large-scale community phenology models (Melaas et al., 2016), and forecast long-term phenology trends (Jeong et al., 2013). Large-scale datasets from China and Europe have already contributed considerably to phenological research (Xu and Chen, 2013; Olsson and Jönsson, 2014; Basler, 2016; Zhang et al., 2017), and the NPN dataset has the potential to meet these needs for North American plant species and communities. However, the features that allow citizen science projects to collect data at large scales can also introduce spatial biases toward cities and easily-accessible areas, and variation in sampling effort and observer skill (Dickinson et al., 2010). With hundreds of participants across North America, the potential for variation among observers in their determination of species identification and dating of phenological events is high. While volunteers have been shown to be accurate at distinguishing different leaf and flower stages for plants, (Fuccillo et al., 2015), observations are sometimes made sporadically across seasons, years, and locations. This means that the quanity and quality of data at a specific site will typically be more variable for citizen science efforts than for intensive, long-term studies.

In order to accurately model and forecast phenology, it is important to understand how the strengths and weaknesses of intensive local studies and large-scale citizen science projects influence both our inferences about biological processes driving phenology and our ability to predict future phenology events. Here, we fit a suite of plant phenology models for the budburst and first-flowering phenophases of 24 plant species to data from both the NPN and a set of intensive long-term studies from the Long Term Ecological Research (LTER) network. We compare the resulting models based on both inference about models and parameters and predictions for unobserved events. We then use this comparison to assess the best methods for both local- and large-scale phenology modeling and to point the way forward for integrating large-scale and local-scale data to determine the best possible models across scales.

## Methods

### Datasets

The National Phenology Network protocol uses status-based monitoring, where via a phone app or web based interface observers answer ‘yes,’ ‘no,’ or ‘unsure’ when asked if an individual plant has a specific phenophase present (Denny et al., 2014). Phenophases refer to specific phases in the annual cycle of a plant, such as the presence of emerging leaves, flowers, fruit, or senescing leaves. Sites in the NPN datasets are located across the U.S. and generally clustered around populated areas (Fig. 1). To represent long-term, intensive phenology studies we used four datasets from North America representing three major ecosystem types (Table 1, Fig. 1). All four long-term studies are located in the U.S. and are part of the Long Term Ecological Research network (LTER). The Harvard Forest and Hubbard Brook Long Term Experimental Forest are located in the northeastern U.S. and are dominated by deciduous broadleaf species. The H.J. Andrews Experimental Forest is a coniferous forest in the coastal range of the western U.S. The Jornada Experimental Range is in the Chihuahua desert of the southwestern U.S.

**Figure 1:**
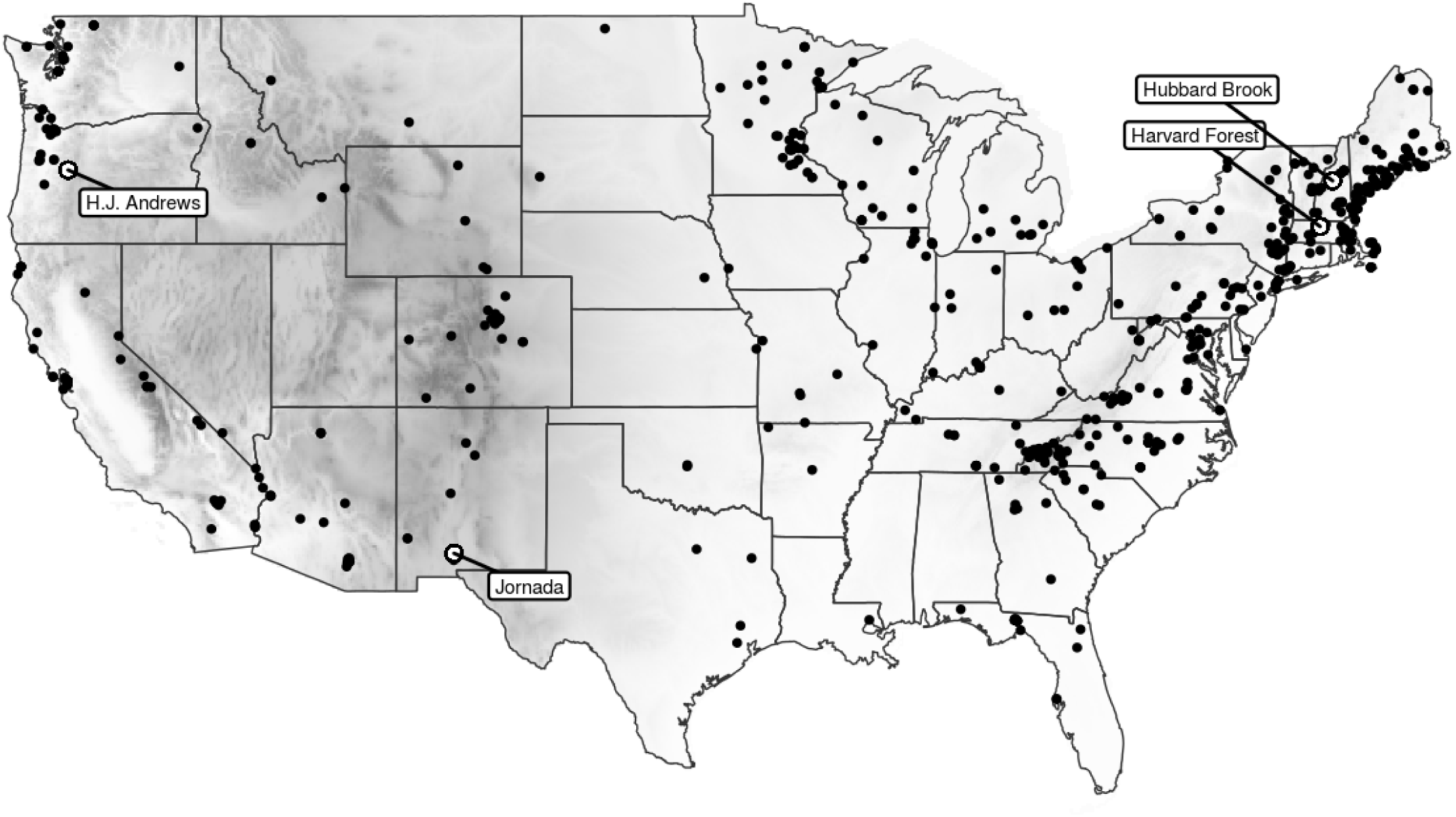
Locations of National Phenology Network sites used (black points) and Long Term Ecological Research sites (labeled circles), with greyscale showing elevation.

**Table 1:**
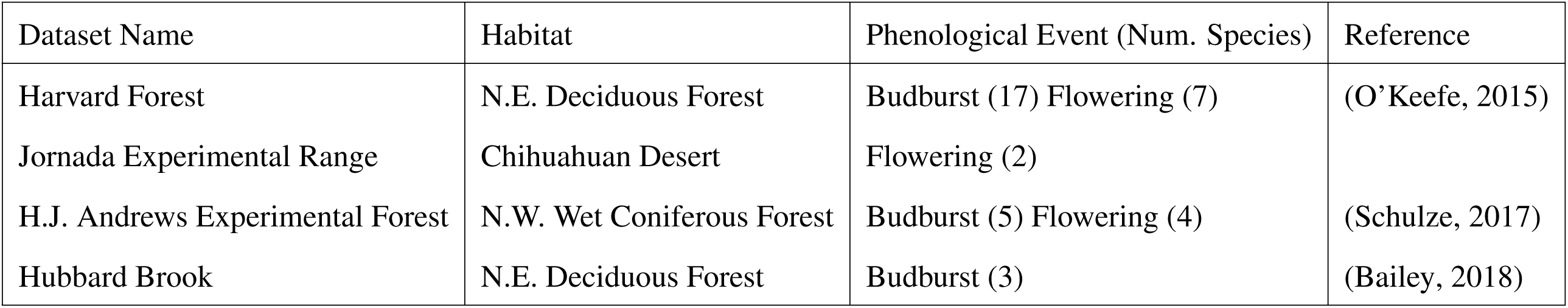
LTER datasets used in the analysis

We downloaded all NPN observations from 2009, when collections began, to 2016 for the following phenophases: Breaking Leaf Buds, Breaking Needle Buds, Emerging Needles, and Open Flowers. The first three phenophases apply to the ‘leaf out’ phase for deciduous broadleafs, evergreen conifers, and pines, respectively. The ‘Open Flowers’ phenophase refers to fully-open flowers and applies to all angiosperms. Hereafter, we will refer to these as either ‘Flowers’ for the Open Flower phenophase, or ‘Budburst’ for all other phenophases. We subset the NPN observations similar to methods outlined in Crimmins et al. (2017). First, ‘yes’ observations for individual plants were kept only if they were preceded by a ‘no’ observation within 30 days. Observations for ‘Budburst’ that were past day of year (DOY) 172, and for ‘Flowers’ that were past DOY 213 were dropped to minimize any influence from outliers. We inferred the observed DOY of each phenophase as the midpoint between each ‘yes’ observation and the preceding ‘no’ observation. Finally, only species that had greater than 30 total observations were kept. Crimmins et al. (2017) only kept observations that were preceded by a ‘no’ within 15 days, and also grouped multiple individuals at single sites to a single observation. We used 30 days to allow for a greater number of species to be compared. We chose not to group multiple individuals at a single site to better incorporate intra-site variability.

In the LTER datasets observation metrics varied widely due to different protocols. To match the NPN data we converted all metrics to binary ‘yes’ and ‘no’ observations for each phenophase (see supplementary methods). As with the NPN data, we inferred the date for each phenophase as the midpoint between the first ‘yes’ observation and most recent ‘no’ observation, and only kept species and phenophases combinations which had at least 30 total observations. After data processing there were 38 species and phenophase combinations (with 24 unique species) common to both the NPN and LTER datasets to use in the analysis (Table 1 & S1).

### Models

It is common to fit multiple plant phenology models to find the one that best represents a specific species and phenophase (Chuine et al., 2013). For each of the 38 species and phenophase combinations in the five datasets (NPN and four LTER datasets), we fit eight phenology models (Table 2). The *Naive model* uses the mean DOY from prior observations as the estimated DOY. The *Linear model* uses a regression with the mean spring (Jan. 1 - March 31) temperature as the independent variable and DOY as the response variable. For the six remaining models, the general form is based on the idea that a phenological event will occur once sufficient thermal forcing units, *F*^*\**^, accumulate from a particular start day of the year (*t*_1_). Forcing units are a transformation of the daily mean temperature. The start day can either be estimated or fixed. For the *Growing Degree Day (GDD)* model, forcing units are the total degrees above the threshold *T*. The *Fixed GDD model* uses the same form but has fixed values for start day (*t*_1_ = Jan 1) and temperature threshold (*T* = 0^*°*^C). The *Alternating model* has a variable number of required forcing units defined as a function of the total number of days below 0^*°*^C since Jan. 1 (*NCD*). The *Uniforc model* is like the GDD model but with the forcing units transformed via a sigmoid function (Chuine, 2000). These models are some of the most commonly used in phenology research and serve as a suitable baseline for comparing the NPN and LTER datasets.

**Table 2:** Phenology models used in the analysis

| Name | DOY Estimator | Forcing Equations | Total Parameters | Reference |
| --- | --- | --- | --- | --- |
| Naive | $\overline{DOY}$ | - | 1 | - |
| Fixed GDD | $\sum_{t=0}^{DOY} R_f(T_i) \geq F^*$ | $R_f(T_i) = \min(T_i, 0)$ | 1 | - |
| Linear | $DOY = \beta_1 + \beta_2 T_{mean}$ | - | 2 | - |
| GDD | $\sum_{t=t_1}^{DOY} R_f(T_i) \geq F^*$ | $R_f(T_i) = \max(T_i - T^*, 0)$ | 3 | - |
| M1 | $\sum_{t=t_1}^{DOY} R_f(T_i) \geq (\frac{L_i}{24})^k F^*$ | $R_f(T_i) = \max(T_i - T^*, 5)$ | 4 | (Blümel and Chmielewski, 2012) |
| Alternating | $\sum_{t=0}^{DOY} R_f(T_i) \geq a + be^{cNCD(t)}$ | $R_f(T_i) = \max(T_i - 5, 0)$ | 3 | (Cannell and Smith, 1983) |
| MSB | $\sum_{t=0}^{DOY} R_f(T_i) \geq a + be^{cNCD_i} + dT_{mean}$ | $R_f(T_i) = \max(T_i - 5, 0)$ | 4 | (Jeong et al., 2013) |
| Uniforc | $\sum_{t=t_1}^{DOY} R_f(T_i) \geq F^*$ | $R_f(T_i) = \frac{1}{1+e^{b(T_i-c)}}$ | 4 | (Chuine, 2000) |

We also fit two models that attempt to capture spatial variation in phenological requirements. The first spatial model, *M1*, is an extension of the *GDD model* which adds a correction in the required forcing using the photoperiod (*L*) (Blümel and Chmielewski, 2012). The second, the *Macroscale Species-specific Budburst model (MSB)*, uses the mean spring temperature as a linear correction on the total forcing required in the *Alternating model* (Jeong et al., 2013). Since there is little to no spatial variation in the LTER datasets, we fit the two spatial models to data from the NPN only. We compared the resulting parameters, estimates, and errors for the NPN-derived *M1* and *MSB* models to their non-spatial analogs (the *GDD* and *Alternating models*, respectively) for each species and phenophase in the LTER data.

We extracted corresponding daily mean temperature for all NPN and LTER observations from the gridded PRISM dataset (PRISM Climate Group, 2004). We parameterized all models using differential evolution to minimize the root mean square error (RMSE) of the estimated DOY of the phenological event. Differential evolution is a global optimization algorithm which uses a population of randomly initialized models to find the set of parameters that minimize the RMSE (Storn and Price, 1997). Confidence intervals for parameters were obtained by bootstrapping, in which individual models were re-fit 250 times using a random sample, with replacement, of the data. We made predictions by taking the mean DOY estimated from the 250 bootstrapped iterations. A random subset consisting of 20% of observations from each species and phenophase combination was held out from model fitting for later evaluation.

### Analysis

As described above, we fit two sets of models for each species and phenophase: one set of models parameterized using only NPN data, and one set parameterized using only LTER data (with the exception of the *M1* and *MSB* models, see above). To compare the inferences about process made by the two datasets, we compared the distribution of each parameter between LTER and NPN-derived models for each species and phenophase combination. Using the mean value of each bootstrapped parameter, we also calculated the coefficient of determination (*R*^2^) between LTER and NPN-derived models among the 38 species-phenophases. In three cases where a species phenophase combination occurred in two LTER sites (Budburst for *Acer saccharum, Betula alleghaniensis*, and *Fagus grandifolia* in the Harvard and Hubbard Brook datasets) they were compared separately to the NPN data.

Models with different parameter values, and even entirely different structures, can produce similar estimates for the date of phenological events (Basler, 2016). Therefore, to compare the predictions and potential forecasts for models fit to the different datasets, we compared the estimated DOY predicted by the LTER and NPN derived models for all held out observations. For each of the eight models, we calculated the coefficient of determination (*R*^2^) between LTER and NPN-derived estimates for estimates made at the four LTER sites and across all NPN sites.

We also directly evaluated model performance using four combinations of models and observed data: A) LTER-derived models predicting LTER observations, B) NPN-derived models predicting LTER observations, C) LTER-derived models predicting NPN observations, and D) NPN-derived models predicting NPN observations. Within each of these scenarios, we calculated the RMSE of the held-out observations for each species, phenophase, and model type. Using the RMSE values, we calculated two different metrics to compare the performance of LTER and NPN-derived models on different data types. The first metric focuses on local-scale prediction by comparing the fits of LTER and NPN-derived models on LTER observations: *RMSE*_*A*_ *-RMSE*_*B*_. The second metric focuses on large-scale prediction by comparing the fits of LTER and NPN-derived models on the NPN data: *RMSE*_*C*_ *-RMSE*_*D*_. These metrics were calculated for each of the model types and 38 species-phenophase combinations. Negative values indicate that LTER-derived models perform better, while positive values indicate that the NPN-derived model performed better. In the three cases where the same species and phenophase combination occurred in two LTER sites, we made the LTER-LTER comparison (scenario A) within each site, not across sites, to focus on local scale prediction when LTER data is available. Absolute RMSE values are provided in the supplement (Fig. S1-S3).

We performed all analysis using both the R and Python programming languages (R Core Team, 2017; Python Software Foundation, 2018). Primary R packages used in the analysis included dplyr (Wickham et al., 2017), tidyr (Wickham and Henry, 2018), ggplot2 (Wickham, 2016), lubridate (Grolemund and Wickham, 2011), prism (Hart and Bell, 2015), raster (Hijmans, 2017), and sp (Pebesma and Bivand, 2005). Primary Python packages included SciPy (Jones et al., 2001), NumPy (Oliphant, 2006), Pandas (McKinney, 2010), and MPI for Python (Dalcin et al., 2011). Code to fully reproduce this analysis is available on GitHub (https://github.com/sdtaylor/phenology_dataset_study and archived on Zenodo (https://doi.org/10.5281/zenodo.1256705)

## Results

The best matches between parameter estimates based on NPN and LTER data were the *Fixed GDD model* (*R*^2^ = 0.49) and the *Linear model* (*R*^2^ = 0.39 for *β*_1_ and −0.05 for *β*_2_). The parameters for all other models had *R*^2^ values <0 indicating that the relationship was worse than no relationship between the parameters (but with matching mean parameter values across the two sets of models) (Fig. 2). The *Naive model* showed a distinct late bias in mean DOY estimates for phenological events, likely resulting from the LTER datasets being mostly in the northern United States compared to the site locations of the NPN dataset (Fig. 2). The large outlier for the *Fixed GDD model* is *Larrea tridentata*; this species’ flower phenology is largely driven by precipitation, which is not considered in the Fixed GDD model (Beatley, 1974). While the *Fixed GDD* and *Linear* models showed reasonable correspondence between parameter estimates, all parameters for individual species and phenophase combinations had different distributions between NPN and LTER-derived models (Fig. S6-S7).

**Figure 2:**
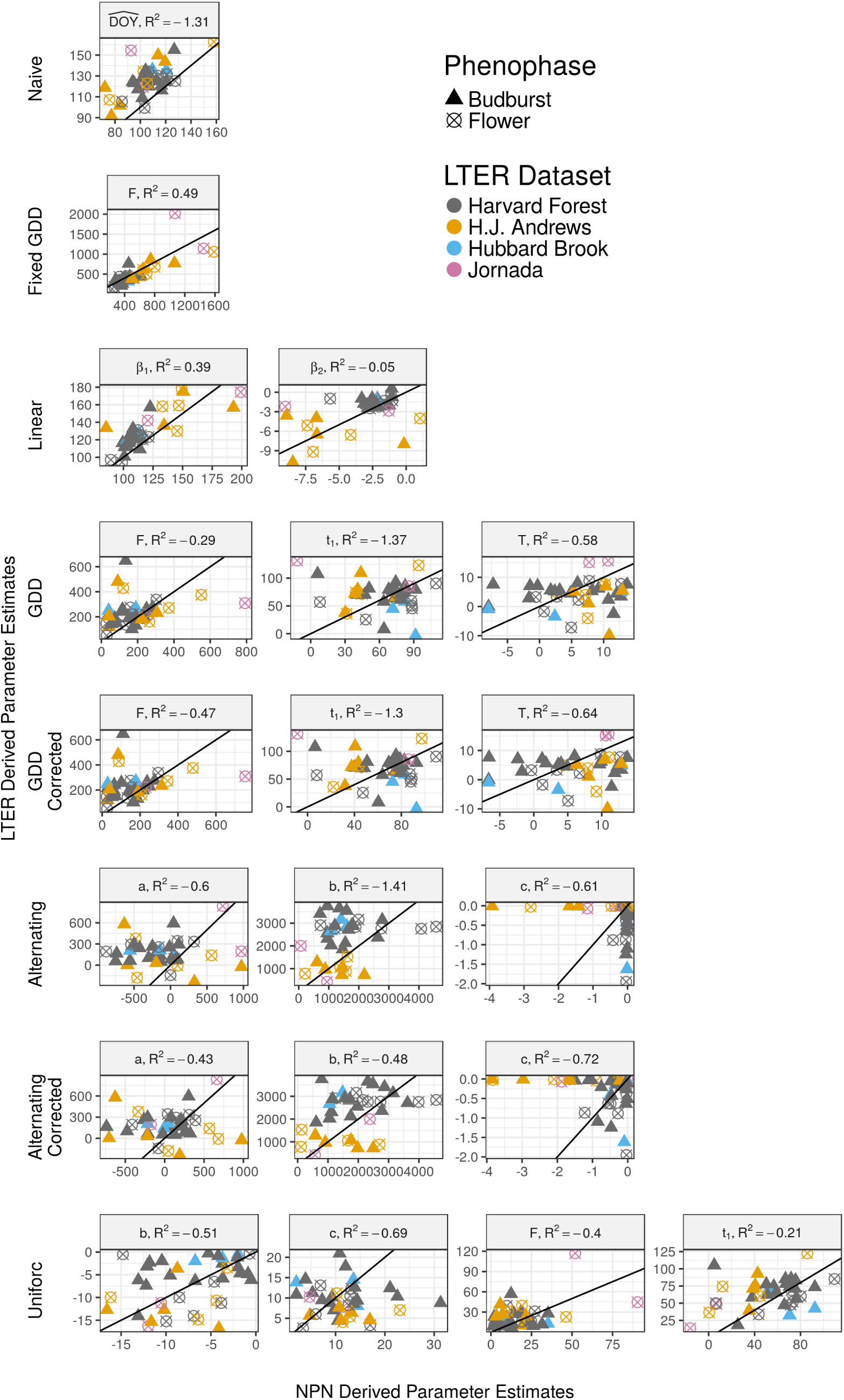
Comparisons of parameter estimates between NPN and LTER derived models. Each point represents a parameter value for a specific species and phenophase, and is the mean value from 250 bootstrap iterations. The black line is the 1:1 line. The *R*^2^ is the coefficient of determination, which can be negative if the relationship between the two parameter sets is worse than no relationship but with the same mean values.

When comparing estimates of phenological events between the two sets of models, many NPN and LTER models produced similar estimates (Fig. 3). The *Fixed GDD model* had the highest correlation between the two models sets at NPN sites (*R*^2^ = 0.82), while the *GDD, M1*, and *Uniforc* models had the highest correlation at LTER sites (*R*^2^ = 0.51, 0.52, and 0.51, respectively). Comparing models with spatial corrections to the non-spatial alternatives, the *MSB* (an extension of the *Alternating model* with a spatial correction based on mean spring temperature, see Table 2 and Methods) improved the correlation between the two datasets over the *Alternating model*. The *MSB model* improved the *R*^2^ from 0.36 to 0.45 at LTER sites, and from −0.23 to −0.15 at NPN sites. The *M1 model* (an extension of the *GDD model* with a spatial correction based on day length) improved the correlation over the *GDD model* only slightly at LTER sites (from 0.51 to 0.52) and did not improve the correlation at NPN sites.

**Figure 3:**
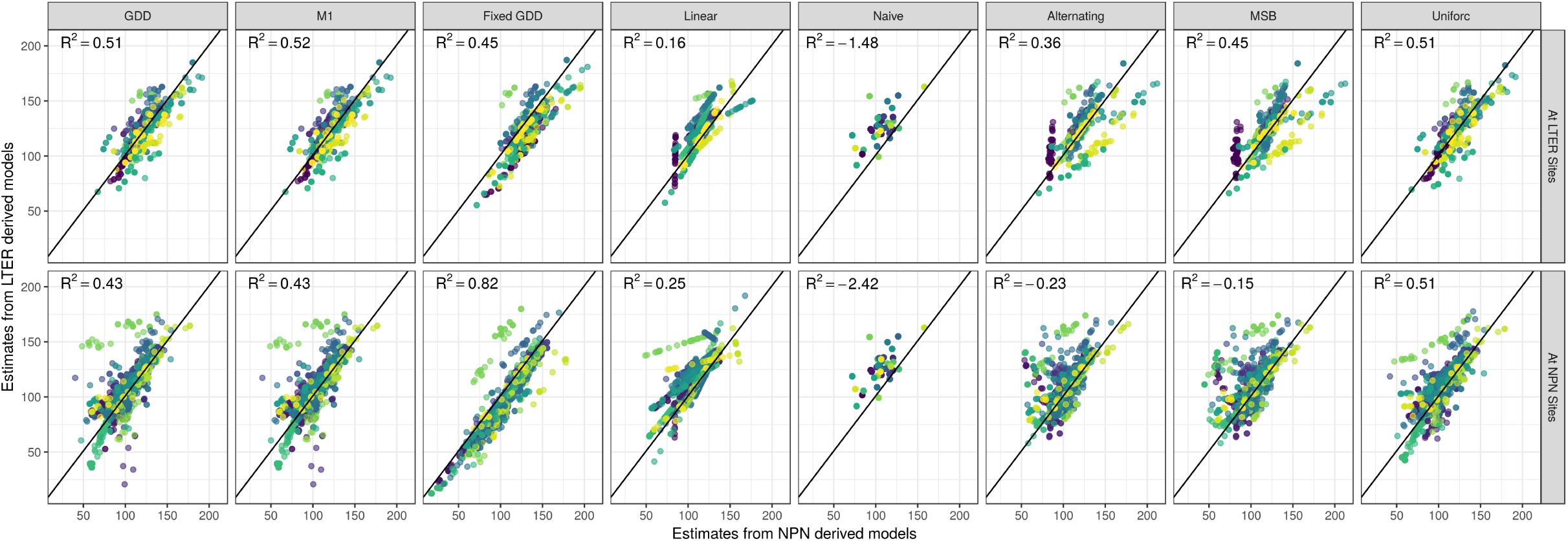
Comparison of predicted day of year (DOY) of all phenological events between NPN and LTER-derived models. Top panels show comparisons at LTER sites and bottom panels show comparisons at NPN sites. Each point is an estimate for a single held-out observation. Colors indicate observations for a single species and phenophase combination.

When comparing the prediction accuracy on held-out data, NPN-derived models made more accurate predictions for held-out NPN observations, and LTER-derived models performed better on held-out LTER observations (Fig. 4). The *Naive* and *Linear* models had the largest differences between the two model sets, while the *Fixed GDD model* had relatively similar errors when evaluated on both NPN and LTER held-out observations. Although the *Fixed GDD model* had the highest agreement in accuracy between NPN and LTER-derived models, it was not the best performing model overall. The *GDD* and *Uniforc* models commonly made the best predictions, having the lowest RMSE in 23% and 40% of cases among NPN-derived models, and 42% and 32% of cases among LTER-derived models, respectively (Fig. S1 & S2).

**Figure 4:**
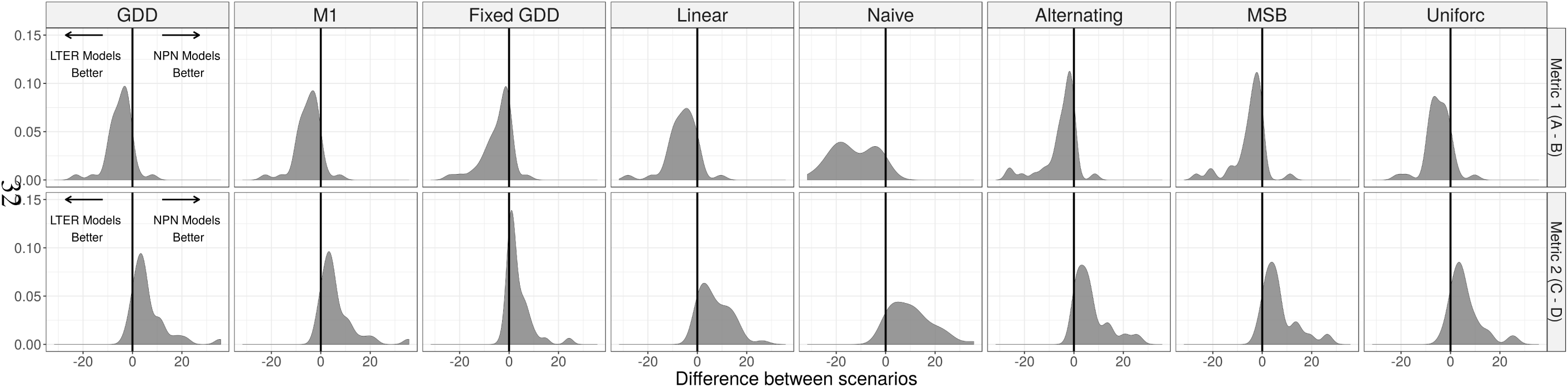
Differences in prediction error between NPN and LTER-derived models. Density plots for comparisons of predictions on LTER data (top row) and NPN data (bottom row). Each plot represents the difference between the RMSE for LTER-derived model and the NPN-derived model, meaning that values less than zero indicate more accurate prediction by LTER-derived models and values greater than zero indicate more accurate prediction by NPN-derived models. Differences are calculated pairwise for the 38 species/phenophase comparisons.

## Discussion

Data used to build phenology models typically falls into two categories: intensive long-term data with long time-series at a small number of locations (e.g., LTER data in this study), and large-scale data with less intensive sampling at hundreds of locations (e.g., NPN data) (Table 3). This data scenario–a small amount of intensive data and a large amount of less intensive data–is common in many areas of science and makes it necessary to understand how to choose between, or combine, data sources (Hanks et al., 2011). We explored this issue for phenology modeling in relation to making predictions and inferring process from models. For inference we found that models based on different data sources resulted in different parameter estimates for all but the simplest models. For prediction we found that models fit to different data sources tended to make similar predictions, but that models better predicted out-of-sample data from the data type to which they were fit. These results are consistent with other research showing that phenology model performance decreases when transferring single-site models to other locations (García-Mozo et al., 2008; Xu and Chen, 2013; Basler, 2016), and with the call for models that better incorporate spatial variation in phenology requirements (Richardson et al., 2013; Chuine and Régnière, 2017). Understanding and making predictions for the phenology of a single location is best served by intensive local-scale data, when available, but large-scale datasets work better for extrapolating phenology predictions across a species range. Thus, the best choice of both data and models depends on the desired research goals.

**Table 3:**
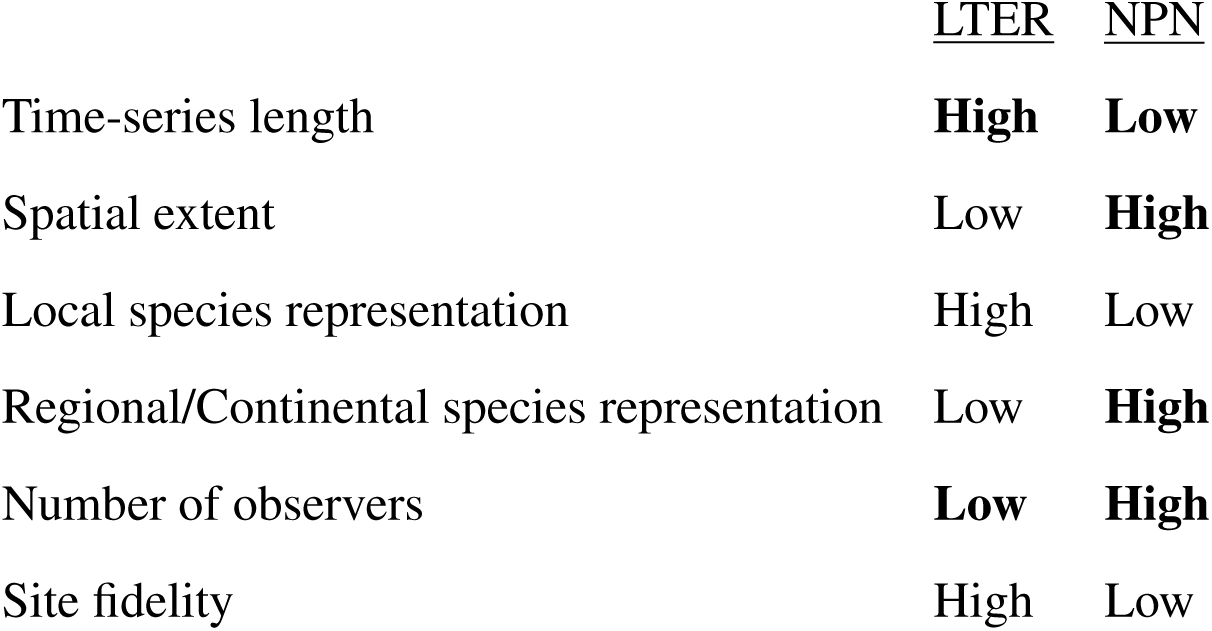
Attributes of the two datasets used in this study. Bold text indicates an attribute is expected to increase over time.

In this study, parameter estimates differed widely within the same phenology model when fit to the two different types of data, except for the simplest process-oriented model: the Fixed GDD (Fig. 2). These differences may be caused by a variety of factors that have different implications for interpreting process-oriented models and their parameters. First, the differences could result from limitations in the sampling of the NPN dataset, leading to less accurate parameter estimates. If this is the case, it would suggest that using LTER data is ideal for making inferences about plant physiology, and that focusing on the Fixed GDD model is best for making inferences when NPN data is all that is available. Second, spatial variation in phenology requirements could drive these differences, because NPN data integrates over that spatial variation, while LTER data only estimates the phenological requirements for a specific site. In this case, NPN data would provide a better estimate of the general phenological requirements of a species, but LTER data would provide a more accurate understanding for a single site. The best solution to this issue would be the development of models that accurately incorporate spatial variation, such as including genetic variation between different populations (Chuine and Régnière, 2017). Third, these differences could result from issues with model identifiability. Since different parameter values can yield nearly identical estimates of phenological events, parameter estimates can differ between datasets even when the underlying processes generating the data are the same. Information about which of these issues may be causing the differences between datasets can be explored using these analyses, as will be explained below.

Despite substantial differences in parameter estimates, LTER and NPN-derived models produced similar estimates for phenological events in most cases (Fig. 3). This greater correspondence between predictions than parameters suggests that more complex models may have identifiability issues. For example, two GDD models with parameters of *t*_1_=1, *F*=10, *T*^*\**^=0 and *t*_1_=5, *F*=5, *T*^*\**^=0 produce nearly identical estimates in many scenarios. This possibility is supported by the fact that the highest correlation between parameter estimates is seen in models with only 1 or 2 parameters. In addition, bootstrap results for more complex models suggest a high degree of variability in parameter estimates and potentially multiple local optima in fits to both NPN and LTER data (Fig. S6-S7). Finally, parameter estimates of more complex models are also not consistent among models for the same species when comparing multiple LTER datasets (Fig. S4-S5). These results are consistent with research showing that models estimating the starting day of warming accumulation from budbreak time-series failed to accurately infer the internal phenology described in the models (Chuine et al., 2016). Basler (2016) suggests that the key component in phenology models is the thermal forcing, with additional parameters being sensitive to over-fitting. Here, our simplest model, the *Fixed GDD model* which uses only a warming component, had the highest correlation among parameters between LTER and NPN datasets. In combination with this previous research, our results warrant caution in interpreting parameter estimates from complex phenology models regardless of the data source used for fitting the models.

While more complex phenology models appear to have identifiability issues, there is also evidence that they capture useful information, beyond the *Fixed GDD model*, based on their ability to make out-of-sample predictions. Based on the RMSE, the *GDD* and *Uniforc* models produce the best out-of-sample predictions for the majority of species and phenophases at both NPN and LTER datasets (Fig. S1 & S2). This demonstrates that the more complex models are capturing additional information about phenology, and that some of the differences between datasets result from differences in either the scales or the sampling of the data. Spatial variation in phenological requirements is known to exist in plants (Zhang et al., 2017). In combination with our results showing observed differences in parameter estimates between LTER sites (Fig. S4-S5), this suggests that variation in phenological requirements across the the range is likely important. However, the models that attempted to address this by incorporating spatial variation did not yield improvements over their base models in our analyses. Specifically, correspondence between parameter estimates (Fig. 2), estimates of phenological events (Fig. 3), and out-of-sample error rates (Fig. 4) for the MSB and *M1* models were essentially the same as the *Alternating* and *GDD* models, respectively. This lack of improvement from incorporating spatial variation could be caused either by models not adequately capturing the process driving the spatial variation, the NPN dataset having biases from variation in sampling effort and/or spatial auto-correlation, or some combination of these factors. Basler (2016) used the *M1* model to predict budburst for six species across Europe and found it was generally among the best models in terms of RMSE, albeit never by more than a single day. Their result was strengthened by having a 40-year time-series across a large region. Chuine and Régnière (2017) listed the incorporation of spatial variation in warming requirements in models as a primary issue in future phenology research. Large-scale phenology datasets, like NPN, will be key in addressing this and other phenological research needs.

In conclusion, our results suggest that both LTER and NPN data provide valuable information on plant phenology. Models built using both data sources yield effective predictions for phenological events, but parameter estimates from the two data sources differ and models from each source best predict that data source’s phenology events. The primary difference in the datasets is spatial scale, but due to trade-offs in data collection efforts, the larger scale NPN data has shorter time-series, less site fidelity and other differences from the intensively collected LTER data (Table 3). These differences can be strengths or potential limitations. Observers sampling opportunistically allows the NPN dataset to have a large spatial scale, but also leads to low site fidelity which limits the ability to measure long-term trends at local scales (Gerst et al., 2016). Tracking long-term trends is the major strength of LTER data, but having a relatively small species pool limits its use in species-level predictive modeling. Due to these differences, the best data source for making predictions depends on the scale at which the predictions are being made. Identifying the most effective data sources for different types and scales of analysis is a useful first step, but the ultimate solution to working with diverse data types is to focus on integrating all types of data into analyses and forecasts (Hanks et al., 2011; Melaas et al., 2016). Our results suggest that methods that can learn from the intensive information available in LTER data in regions where it is available, and simultaneously use large-scale data to capture spatial variation in phenological requirements will help improve our ability to understand and predict phenology. Data integration efforts should also leverage data from remote sensing sources such as the PHENOCAM network or satellite imagery, which have both a large spatial extent and high temporal resolution (Richardson et al., 2018). Data integration provides the potential to use data from many sources to produce the best opportunity for accurate inference about, and forecasting of, the timing of biological events.

## Acknowledgments

This research was supported by the Gordon and Betty Moore Foundation’s Data-Driven Discovery Initiative through Grant GBMF4563 to E.P. White. We thank the developers and providers of the data that made this research possible including: the USA National Phenology Network and the many participants who contribute to its Nature’s Notebook program; the H.J. Andrews Experimental Forest research program, funded by the National Science Foundation’s (NSF) LTER Program (DEB-1440409), US Forest Service Pacific Northwest Research Station, and Oregon State University; the Jornada Basin LTER project (NSF Grant DEB-1235828); the Hubbard Brook Experimental Forest, which is operated and maintained by the USDA Forest Service, Northern Research Station, Newtown Square, PA.; the Harvard Forest LTER; and the PRISM Climate Group at Oregon State University.

